# Is there a right place? The effect of oviposition site selection on offspring survival in a glassfrog

**DOI:** 10.1101/2024.08.18.608513

**Authors:** Francesca N. Angiolani-Larrea, Anyelet Valencia-Aguilar, Marina Garrido-Priego, Mélissa Peignier, Jaime Culebras, Lelis Jindiachi, José G. Tinajero-Romero, Juan M. Guayasamin, Eva Ringler

**Affiliations:** Division of Behavioural Ecology, Institute of Ecology and Evolution, University of Bern, Bern, Switzerland; Institute of Animal Physiology, Department of Animal Physiology and Molecular Biomedicine, Justus-Liebig-University Giessen, Giessen, Germany; Photo Wildlife Tours, Quito, Ecuador; Fundación Cóndor Andino, Quito, Ecuador; Pueblo Shuar Arutam, Federación Interprovincial Centros Shuar (FICSH), Sucúa, Ecuador; Escuela de Biologia, Universidad de Costa Rica, San José, Costa Rica; Universidad San Francisco de Quito USFQ, Laboratorio de Biología Evolutiva, Colegio de Ciencias Biológicas y Ambientales COCIBA, Instituto BIÓSFERA-USFQ, Campus Cumbayá, Quito 170901, Ecuador; Tandayapa Cloud Forest Station, Universidad San Francisco de Quito USFQ, P.O. Box 17-1200-841, Quito, Ecuador

## Abstract

The choice of where to breed can have fundamental consequences for offspring development and survival. In amphibians, desiccation is one of the biggest threats to survival, especially in species that deposit their clutches in terrestrial habitats. In several species, hydration of the clutch is ensured by a caregiving parent, but in species without extended care, it might be the selection of a suitable oviposition location that helps secure a constant external source of hydration. We used *Teratohyla spinosa*, a Neotropical glassfrog without extended parental care, to test the effect of oviposition site selection on offspring development and survival. Previous observations have revealed that this species preferentially deposits eggs on the underside of leaves close to their margins. We hypothesized that *T. spinosa* strategically chooses this position on the leaves to ensure clutch hydration during embryonic development. To this end, we performed a clutch translocation experiment where we manipulated the location of clutches and compared their level of hydration, hatching time, and mortality rate to control clutches. We found that clutch hydration and mortality were not affected by the location on the leaf. These findings suggest that the oviposition site selection in *T. spinosa* does not affect clutch survival – at least under high humidity conditions.

## Introduction

In many animal species, the choice of where to breed and to place offspring can have a major impact on offspring survival, development, and reproductive potential [1,2]. Individuals can maximize their reproductive success by trading off the allocation of resources to their current or future offspring, and their own needs [1], though these strategies are not necessarily mutually exclusive. In many species, the choice of where to breed is often non-random and influenced by factors such as intraspecific competition, habitat structure, predation pressure, and climatic variables [2].

So far, breeding site selection has mostly been studied at a macro environmental level. It has been shown that many species exhibit preferences of where to breed [3,4] with respect to vegetation (e.g.: [5]) and water availability (e.g.: [6]), habitat composition, predators, or conspecific and parasite densities (e.g.: [7,8]). However, in many cases, the choice of the breeding site can be influenced by even more fine-scale parameters [3,9], such as microhabitat composition, local topography, or edge effects. For example, in two sympatric species of polar seals it was found that both use drifting ice to breed; while one species was less choosy about the specific breeding spot, the other one was highly selective about the particular topographical characteristics of the breeding site [10]. Moreover, in some species of tree frogs (Hylidae) water depth, distance to the pond, temperature, and pond size play a key role when choosing their breeding sites [11]. The analysis of factors across different scales therefore can inform about the selective pressures that have led to differences in reproductive behaviours across and within species.

In many amphibian species, it has been shown that oviposition site selection is key for offspring survival and development (e.g.: [5,7,12–15]). Desiccation is one of the biggest threats to egg survival, especially in species that deposit their clutches outside of large water bodies [5,16–19]. As no hard layers protect their eggs [20], female frogs are expected to select oviposition locations that ensure a constant source of hydration [6,12,16,21].

Glassfrogs (family Centrolenidae) are a good model for investigating the factors that shape oviposition site selection. These Neotropical arboreal frogs deposit their clutches in substrates neighbouring or overhanging streams and are characterized by their transparent and translucent skin [22]. Many species tend to have preferential locations for oviposition at exact positioning on vegetation or other substrates, such as moss [23], rocks in the splash area of waterfalls, the tip of leaves, or the upper or under side of leaves [24]. In glassfrogs, several species exhibit extended male or female uniparental care, which is expressed in form of guarding, brooding and defence against predators [19,25–28]. Such extended forms of care did not evolve in other glassfrog species [25,27], which either leave the oviposition site shortly after mating [23] or show short-term brooding of the clutch [24,27]. Therefore, the selection of a suitable oviposition location in these species may secure constant external source of hydration.

We investigated the possible benefits of preferential locations for oviposition in the species *T. spinosa.* This species lays the eggs on the underside of leaves close to their margins [24]. We hypothesized that the oviposition location would facilitate the hydration of eggs and thereby reduce their mortality risk. In the rainforest, water in the form of airborne humidity and rain usually contacts the surface of vegetation before evaporating or falling to the ground. When leaf surfaces are saturated with water, fused drops will slide from the edges to the tip of the leaves before dripping. We experimentally tested whether the position of the clutch on the leaf affects its hydration level and survival, and whether the position of the clutch on the leaf is important for embryo development and survival.

## Materials and Methods

### Study site

Our study took place in Canandé Reserve, Esmeraldas province, Ecuador, along a 500m transect throughout the length of a creek (0° 31’ 24.7’’ N, 79° 12’ 45.6’’ W). The reserve is located between two Biodiversity hotspots: Tumbes-Chocó-Magdalena and Tropical Andes [29], within the Lowland Evergreen Forest ecosystem [30]. The site is characterized by tropical humid seasonal weather and a rainy season that takes place from November to May. The creek floor and margins are heterogeneous, changing from flat boulders to clay. The vegetation corresponds to secondary forest, with old canopy trees higher than 10m and herbaceous plants densely covering the margins of the creek (Fig. 1).

**Figure 1.**
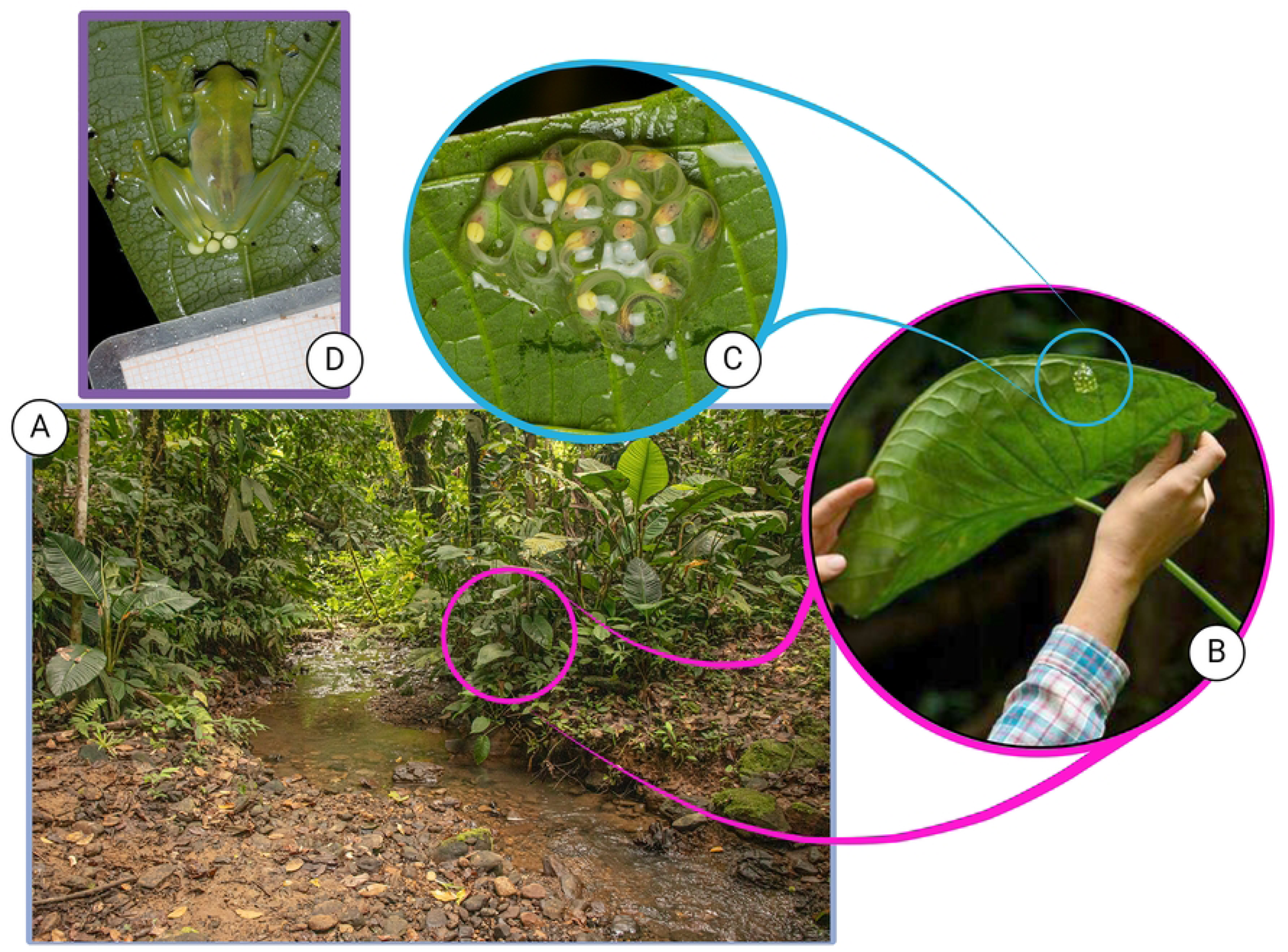
Study system. A) Sampling was done throughout 500m of a creek surrounded by secondary forest, B) Clutches of *Teratohyla spinosa* as naturally occurring. Placed on the underside of leaves near the margins, C) Detail of an old clutch, D) Mother of *T. spinosa* performing brooding after oviposition. Photos A by FNAL and B-D by JC. Created with BioRender.com

### Study species

*Teratohyla spinosa* (Centrolenidae) is a small (<2.5 cm), nocturnal glassfrog, with most of the adult activity taking place in the vegetation alongside rivers, small streams, and creeks [24]. The reproductive activity of *T. spinosa* occurs mainly during the rainy season, when males vocalize to attract mates. Females approach males to engage in amplexus for several hours. During this time, the amplectant pair will hop around vegetation in the close surroundings until selecting a leaf. The female will deposit a jelly-rich clutch of eggs on the underside of the selected leaf, close to the edges or tip (Fig 2A, [24]). Neither males nor females of *T. spinosa* show extended parental care [31].

**Figure 2.**
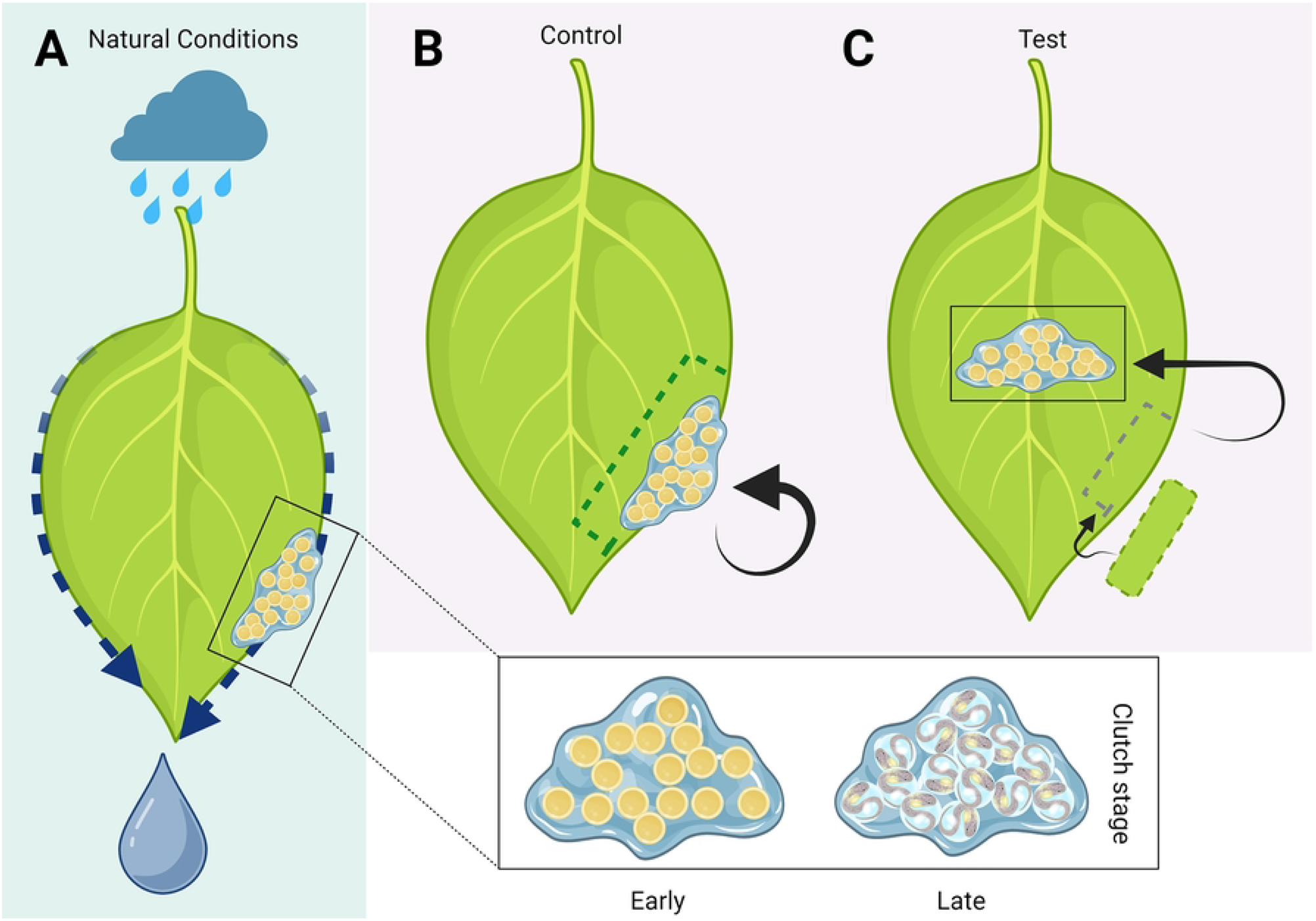
Experimental design to test the effect of clutch position and embryonic survivorship. A) Natural conditions, with the egg clutch located on the edge of the underside of leave. In the box are depicted the developmental stages of the clutches classified as early or late (see Methods). B) Control clutches: on the same position as in natural conditions. C) Test clutches: egg clutches translocated to the centre of the underside of leaves, avoiding contact with the edges. The hole left by the cut area was patched with a piece of leaf from the same plant. Created with BioRender.com

### Field work – experiment

The experiment was carried out from April 16^th^ to June 11^th^, 2022. Clutches were randomly assigned to the experimental or control condition. For the control condition, we cut out the area around the clutch and sewed it with thread and needle back to the original location, to ensure similar handling procedures across conditions (Fig 2B). For the experimental condition, we cut out the clutch in the same way as for the control condition, but the clutches were sewn in the centre of the leaf, aiming for the maximum distance away from the edges. To increase stability of all attachments, we additionally sewed another piece of leaf from the same plant on top of the reattached piece of the test and control clutches. To maintain the shape of all leaves in the test condition, we patched a piece of leave to the removed area (Fig 2C).

All clutches were monitored for hydration, mortality and hatching twice a day, once during the day and once at night, until all tadpoles of the clutch had hatched (mean number of tadpoles = 11,6; SD = 1,9) or were lost. We consider mortality as death by desiccation, predation, fungal infection, parasitism, lack of fertilization and unknown causes. During each observation we turned over the leaves and took lateral and zenithal photographs of the clutch with measuring paper. The transect was monitored daily to find new clutches.

### Data collection

For all clutches we assessed the following parameters: level of hydration, which was measured by the thickness (mm) of the clutch at the highest point. We used lateral views of the clutches measuring from the highest point of the clutch to the nearest leaf point in a 90° angle. Area of the clutch (mm^2^), for which we used zenithal photographs using the jelly margins as limits for the area of the leaf occupied by the clutch. The number of eggs alive and dead/unfertilized in each clutch on each observation; age of the clutch counted as days since oviposition; ratio between the area of the clutch divided by the number of eggs it possessed; the stage of development in which the spawn was at the moment it was found, dividing this into *early* stages (from oviposition to Gosner’s stage 19 [32]) and *late* stages; days until full hatching, counting as days from the first tadpole until the last one to hatch; and the width of the leaf (cm) in the thickest part of the leaf. We obtained daily temperature and humidity data from the locality proportioned by the Reassembly project [33], and we calculated the Temperature and Humidity Index (THI) with the formula THI = 0.8*T + RH*(T-14.4) + 46.4, following Mader et al [34] using the average daily values. Additional variables we accounted for are days passed until the clutch reached hatching capacity of the larvae, the number of days spent on the experiment and percentage of mortality. All photos were analysed using ImageJ (version 1.53r).

Our study followed Good Scientific Practice (GSP) guidelines and the ASAB for the ethical treatment of nonhuman animals in behavioural research and teaching [35]. The Jocotoco Research Station and the ‘Ministerio del Ambiente, Agua y Transición Ecológica’ (permit number: MAATE-DBI-CM-2022-0245) granted us working permits.

### Statistical analyses

We conducted all statistical analyses in R v3.6.0 (R Core Team 2020) using RStudio (RStudio Team 2020). We report our results following Muff *et al*. [36].

We used a Bayesian framework to determine what factors influence the thickness, mortality, and development rate of a clutch. We assumed statistical significance if the 95% credible intervals did not overlap 0.

We used a gaussian Bayesian linear mixed model (MCMCglmm package, [37]) to determine the effect that the position of the clutch has on its level of hydration with thickness as response variable. As fixed effects, we added an interaction between the condition (relocated *versus* control) and the days since oviposition, as well as the width of the leaf, the daily THI, and the number of eggs per area. To account for the repeated measures, we also added the clutch ID as random effect. We used a weak prior for one response and one random variable.

We built a Bayesian model with a Poisson distribution (MCMCglmm package, [37]) to determine what factors influence the hatching time. We used days until full hatch as response variable, and as fixed effects the condition, the average THI during the experiment and the average number of eggs per area during the experiment. Because days until full hatch was a non-integer, we multiplied the values by ten to allow the model to run. We used a weak prior for one response variable.

For these two models, we set a number of iterations of 1 000 000, with a burnin of 10 000 and a thinning interval of 100. We verified the absence of autocorrelation (correlation between lags < 0.1), that sufficient mixing was reached (plots of MCMC chains), and that we ran the Markov chain long enough (Heidelberg and Welch diagnostic tests; [37]).

To determine what factors influence the mortality rate of a clutch, we build a zero-inflated beta regression in a Bayesian framework (brms package, [38]. We set the mean, phi and zero-inflated part similarly with the percentage of mortality at the end of the experiment as response variable, and as fixed effects the condition, the average THI during the experiment, the average number of eggs per area during the experiment and the average thickness of the clutch during the experiment. Using the “get_prior” function, we set a specific prior (3, 0, 2.5) for the intercept class elements and a normal prior for the b class elements. We ran this model 2 000 times on 4 chains with a warmup of 1 000. Because beta regression does not allow for ones, we set all ones to 0.999 to allow the model to run. We verified the absence of autocorrelation and sufficient mixing using diagnostic plots.

Additionally, we performed a univariate COX regression [39] to obtain survival probability differences between conditions with position on the leaf as hazard risk. In this case, we assumed statistical significance if the 95% credible intervals did not overlap 1. We used days since oviposition as response variable and censoring status consists of complete clutch death as event and censored clutches include clutches with at least one embryo hatched, successfully hatched clutches, and clutches that did not hatch until end of experiment but were still alive. In this model we are considering the clutch as a unit and do not account for death events of individual embryos.

## Results

We did not find clear evidence that the condition, the days since oviposition, the width of the leaf or the daily THI, were associated to the level of hydration of a clutch (Table 1). There was week indication that the more eggs per area the lower was the level of hydration, but results were marginally significant (Table 1; 95% CI = [−19.90; 0.079], *p-value* = 0.053).

**Table 1.** Summary of confidence intervals and *p* value of MCMC model for level of hydration of clutch.

|  | Lower 95% CI | Upper 95% CI | *p*MCMC |
| --- | --- | --- | --- |
| Intercept | 0.209 | 16.299 | 0.057 |
| Condition | -1.509 | 1.2151 | 0.853 |
| Days since oviposition | -0.130 | 0.010 | 0.100 |
| Width of leaf | -0.020 | 0.056 | 0.336 |
| THI | -0.112 | 0.103 | 0.914 |
| Number of eggs per area | -19.909 | 0.079 | 0.053 |
| Condition*days since oviposition | -0.143 | 0.033 | 0.218 |

We also found no evidence that the condition, the average THI during the experiment, the average number of eggs per area during the experiment and the average thickness of the clutch during the experiment were associated to the mortality rate of the clutch (Table 2).

**Table 2.** Summary of estimates and confidence intervals for zero-inflated beta regression model. S.E.: Standard Error; CI: Confidence Interval.

|  | **Estimate** | **S.E** | **Lower 95% CI** | **Upper 95% CI** |
| --- | --- | --- | --- | --- |
| Intercept | 12.79 | 32.37 | -48.85 | 77.02 |
| $\Phi$ Intercept | 11.12 | 29.92 | -47.67 | 71.51 |
| Zi Intercept | 19.6 | 20.21 | -6.95 | 66.14 |
| Condition | -0.48 | 0.58 | -1.62 | 0.65 |
| THI | -0.15 | 0.44 | -1.02 | 0.69 |
| Number of eggs per area | -0.09 | 1.03 | -2.11 | 1.92 |
| Thickness | -0.15 | 0.22 | -0.6 | 0.28 |
| $\Phi$ Condition | 0.03 | 0.57 | -1.09 | 1.16 |
| $\Phi$ THI | -0.17 | 0.4 | -0.98 | 0.63 |
| $\Phi$ Number of eggs per area | -5.07 | 11.54 | -27.49 | 17.25 |
| $\Phi$ Thickness | 0.06 | 0.24 | -0.39 | 0.55 |
| Zi Condition | 0.84 | 0.83 | -0.69 | 2.56 |
| Zi THI | -0.34 | 0.28 | -0.98 | 0.01 |
| Zi Number of eggs per area | 25.99 | 21.38 | -13.53 | 69.3 |
| zi Thickness | 0.43 | 0.28 | -0.1 | 0.99 |

From the univariate COX regression, we found week evidence of differences in clutch survival between conditions. The probability of death was 62% higher in control clutches, however, this result was not statistically significant (Fig.3, HR = 1.38, 95%; CI = [0.40; 4.78], *p*-value = 0.6).

**Figure 3.**
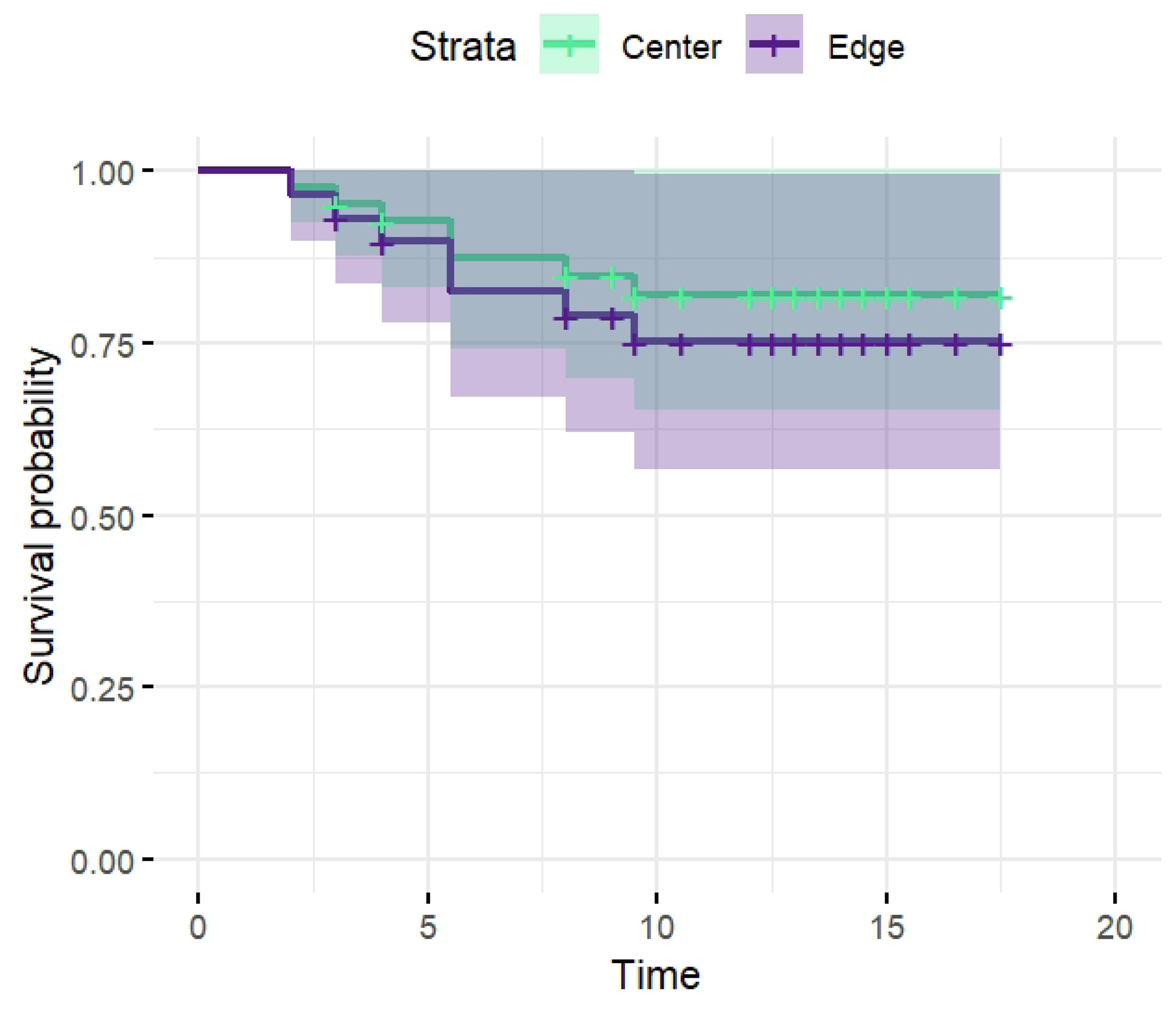
Survival probability of control and experimental clutches of *Teratohyla spinosa*. Probability of survival in both conditions decreased similarly with time (*p* = 0.53)

Lastly, we found no evidence that the condition, THI or eggs per area influenced the number of days it took until full hatch (Table 3).

**Table 3.** Summary of confidence intervals and *p*-value of MCMC model for time of hatch.

| | Lower 95% CI | Upper 95% CI | $p$MCMC |
| --- | --- | --- | --- |
| Intercept | -15.56 | 74.28 | 0.4183 |
| Condition (test-control) | -0.91 | 0.49 | 0.575 |
| THI | -0.97 | 0.25 | 0.227 |
| Eggs per area | -6.82 | 29.98 | 0.206 |

## Discussion

We used a wild population of *T. spinosa* to investigate the effects of oviposition site choice on embryonic development and survival. We hypothesised that the close contact of the clutches with the margins of the leaves offered direct benefits to the eggs in terms of hydration, leading to enhanced development and hatching success. To this end, we manipulated the positioning of clutches on leaves to investigate potential adaptive benefits related to clutch hydration, development, and survivorship.

Contrary to our expectation, we did not find an effect of condition on hydration levels in the clutches (Fig 4). This suggests that hydration is not improved in clutches located at the edge of the leaf. Test and control clutches did not differ in thickness throughout their development. This finding was not what we had expected, given the big threat that desiccation poses to terrestrial/arboreal clutches [40]. In some glassfrog species, parents remain with their eggs almost throughout the entire development, and actively hydrate their eggs (e.g., *Centrolene peristictum* [41], *Hyalinobatrachium aureoguttatum* [42], *Ikakogi tayrona* [43]). Parental removal experiments have shown that the absence of the caring parent leads to a considerable increase in offspring mortality rates [41,44–46]. However, many species of glassfrogs do not exhibit extended parental care and only perform brood once just after oviposition to boost hydration in the jelly-rich clutches [26,27]; although, it is not the case for *Espadarana prosoblepon* [23]. *Teratohyla spinosa* eggs, along with those of other species that lack extended parental care, are embedded in a highly absorbent jelly matrix. It has been proposed for these species that the jelly may be fit for withstanding desiccation under controlled, semi-natural conditions ([31], although in this experiment sample size for *T. spinosa* was one). In our study, we found that under field conditions the embryos do not show signs of dehydration.

**Figure 4.**
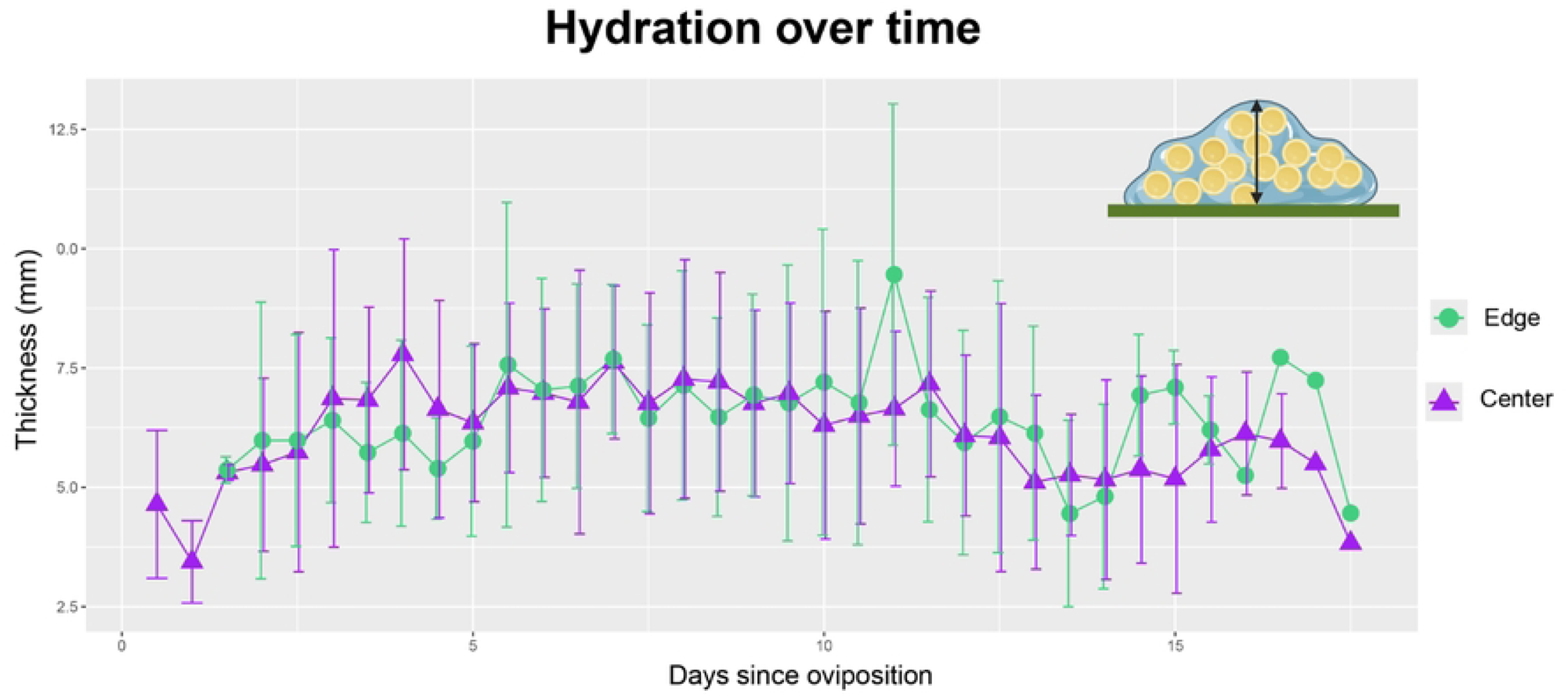
Hydration levels in control and experimental clutches of *Teratohyla spinosa*. Average clutch thickness per day for both experimental conditions: clutches on the edge (control) and on the centre of the leaves (translocated). Created with BioRender.com

Hydric stress has proven to be a driver for changes in developmental rate and hatching plasticity in other species [17,47,48]. We did not observe any signs of hydric stress during development in our study, as test and control clutches did not differ in hatching times. Likewise, there was no indication of hydric stress affecting survivorship, as mortality rates did not differ between conditions. One possible reason for this lack of difference could be that, although we do not measure the weather conditions per oviposition site, these were quite stable overall and humid during the entire experiment, as they were performed during the peak of the rainy season [49]. These conditions might change especially at the beginning and the end of the rainy season when humidity is lower and rainfall is more unpredictable. Eventually, clutches may benefit from increased hydration during periods of low rainfall. Moreover, highly unpredictable weather conditions are more frequent due to climate change, making amphibians more susceptible to extreme weather shifts [16], especially those with terrestrial reproduction [19]. Therefore, future studies should investigate the effects of more variable environmental conditions on clutch development and survival.

Refsnider and Janzen [1] summarized drivers for oviposition site selection into six categories: 1) maximizing embryo survival, 2) maximizing maternal survival, 3) modifying offspring phenotype, 4) proximity to suitable habitat for offspring, 5) maintaining natal philopatry, and 6) indirect oviposition-site choice via mate choice. In our study, we observed no differences to offspring survival or hatching time across conditions. Regarding indirect site-selection via mate choice, we observed males calling from the same areas on consecutive nights, suggesting some form of site fidelity. This indicates active participation by both sexes in oviposition site selection. By choosing the most suitable mate, the female simultaneously selects an oviposition site on a macro scale. However, the decision of the exact location to deposit the eggs at a micro-scale remains unclear. Lastly, there might be benefits for the females associated to the egg laying process itself (i.e. it might be easier for them to lay eggs on the edges), there might also be benefits to the offspring that only manifest at a later stage. These hypotheses remain highly speculative as we did not include such factors in our analysis.

In summary, we found no evidence that clutch placement on the edge of leaves matters for embryonic development and hatching success. We question if an effect might only become apparent at later developmental stages or under more extreme climatic conditions. Alternatively, the placement of clutches at the edges of leaves might reflect some benefit to the female during the egg laying process. Future studies should therefore test for further possible adaptive functions of particular clutch placement across different species.

## Acknowledgements

We want to thank Edith Villa-Galaviz and Santiago Erazo from the REASSEMBLY project for proportioning weather data used in this study. We also thank Fundación Jocotoco, Reserva Canandé, Katrin Krauth, Chiara Correa and Jocotoco’s personnel. We also want to thank Christoph Netz, Max Ringler, Oceane LaLoggia and the Hasli team for their valuable contributions during the development of this project.

## Supporting Information

**S1 Text.** Spanish version of manuscript

